# Dynamic barriers modulate cohesin positioning and genome folding at fixed occupancy

**DOI:** 10.1101/2024.10.08.617113

**Authors:** Hadi Rahmaninejad, Yao Xiao, Maxime M.C. Tortora, Geoffrey Fudenberg

**Affiliations:** Department of Quantitative and Computational Biology, University of Southern California, Los Angeles, USA

## Abstract

In mammalian interphase cells, genomes are folded by cohesin loop extrusion limited by directional CTCF barriers. This interplay leads to the enrichment of cohesin at barriers, isolation between neighboring topologically associating domains, and elevated contact frequency between convergent CTCF barriers across the genome. However, recent *in vivo* measurements present a puzzle: reported residence times for CTCF on chromatin are in the range of a few minutes, while lifetimes for cohesin are much longer. Can the observed features of genome folding result from the action of relatively transient barriers? To address this question, we developed a dynamic barrier model, where CTCF sites switch between bound and unbound states with rates that can be directly compared with biophysical measurements. Using this model, we investigated how barrier dynamics would impact observables for a range of experimental genomic and imaging datasets, including ChIP-seq, Hi-C, and microscopy. We found the interplay of CTCF and cohesin binding timescales influence the strength of each of these features, leaving a signature of barrier dynamics even in the population-averaged snapshots offered by genomic datasets. First, in addition to barrier occupancy, barrier bound times are crucial for instructing features of genome folding. Second, the ratio of boundary to extruder lifetime greatly alters simulated ChIP-seq and simulated Hi-C. Third, large-scale changes in chromosome morphology observed experimentally after increasing extruder lifetime require dynamic barriers. By integrating multiple sources of experimental data, our biophysical model argues that CTCF barrier bound times effectively approach those of cohesin extruder lifetimes. Together, we demonstrate how models that are informed by biophysically measured protein dynamics broaden our understanding of genome folding.

## Introduction

Genome-wide chromosome conformation capture (Hi-C) reveals that mammalian interphase chromosomes are partitioned into a series of topologically associating domains (TADs) (McCord, Kaplan, and Giorgetti 2020). Boundaries between adjacent TADs are largely specified by binding sites for the 11- zinc finger protein CTCF (van Ruiten and Rowland 2021). Boundaries also display enrichment for cohesin by ChIP-seq and are often decorated by punctate dots in Hi-C contact maps between convergent CTCF sites. Acute depletion of CTCF leads to genome-wide loss of TADs and dots in Hi-C contact maps as well as cohesin ChIP-seq enrichment (Nora et al. 2017). Altered CTCF binding even at individual positions can lead to dysregulated gene expression and disease (Lupiáñez et al. 2015).

CTCF is thought to exert its influence on genome folding by acting as a barrier to loop extrusion by the cohesin complex (van Ruiten and Rowland 2021; Fudenberg et al. 2017). During loop extrusion, cohesin binds to chromatin and acts as a motor to progressively create larger chromatin loops until it dissociates. CTCF is thought to act as a directional barrier to impede cohesin extrusion. The initial understanding of the genome-wide consequences of loop extrusion largely relied on comparisons between simulations and static snapshots obtained from genomic data (van Ruiten and Rowland 2021; Fudenberg et al. 2017).

Observations in living cells, however, argue strongly for a dynamic view of genome organization (Hansen 2020). Protein dynamics of CTCF and cohesin have been quantified with Fluorescent Recovery after Photobleaching (FRAP) and Single Particle Tracking (SPT) in a number of cellular contexts (**Fig. 1A, Table S1**). FRAP data indicates cohesins reside on chromatin with lifetimes of 20-30 minutes (Hansen et al. 2017; Gerlich et al. 2006; Tedeschi et al. 2013; Morales et al. 2020) while CTCF resides with shorter 2-11 minute lifetimes (Hansen et al. 2017; Oomen et al. 2019; Soochit et al. 2021; Kieffer-Kwon et al. 2017; Nakahashi et al. 2013; Hansen et al. 2020; Narducci and Hansen 2024). SPT for CTCF indicated a slightly more rapid 1 minute lifetime (Hansen et al. 2017). Tracking DNA, often via integrated arrays of fluorescent reporters, offers a complementary view of dynamics. Importantly, even pairs of genomic loci that appear as salient dots in Hi-C maps are not stably co-localized, and make 5-30 minute contacts (Gabriele et al. 2022; Mach et al. 2022). Single molecule footprinting (SMF), an emerging genomic technology, provides estimates of occupancy of individual CTCF sites. Even strongly bound sites appear to plateau at 70% inferred occupancy (Sönmezer et al. 2021), slightly above an estimated genome-wide average occupancy of 50% (Cattoglio et al. 2019). Each of these technologies argue that dynamic CTCF barriers may enable extruding cohesin motors to bypass individual sites. *In vivo*, however, no existing approach can concomitantly track multiple genomic positions, cohesin, and CTCF and read out their instantaneous occupancy in real time.

**Figure 1.**
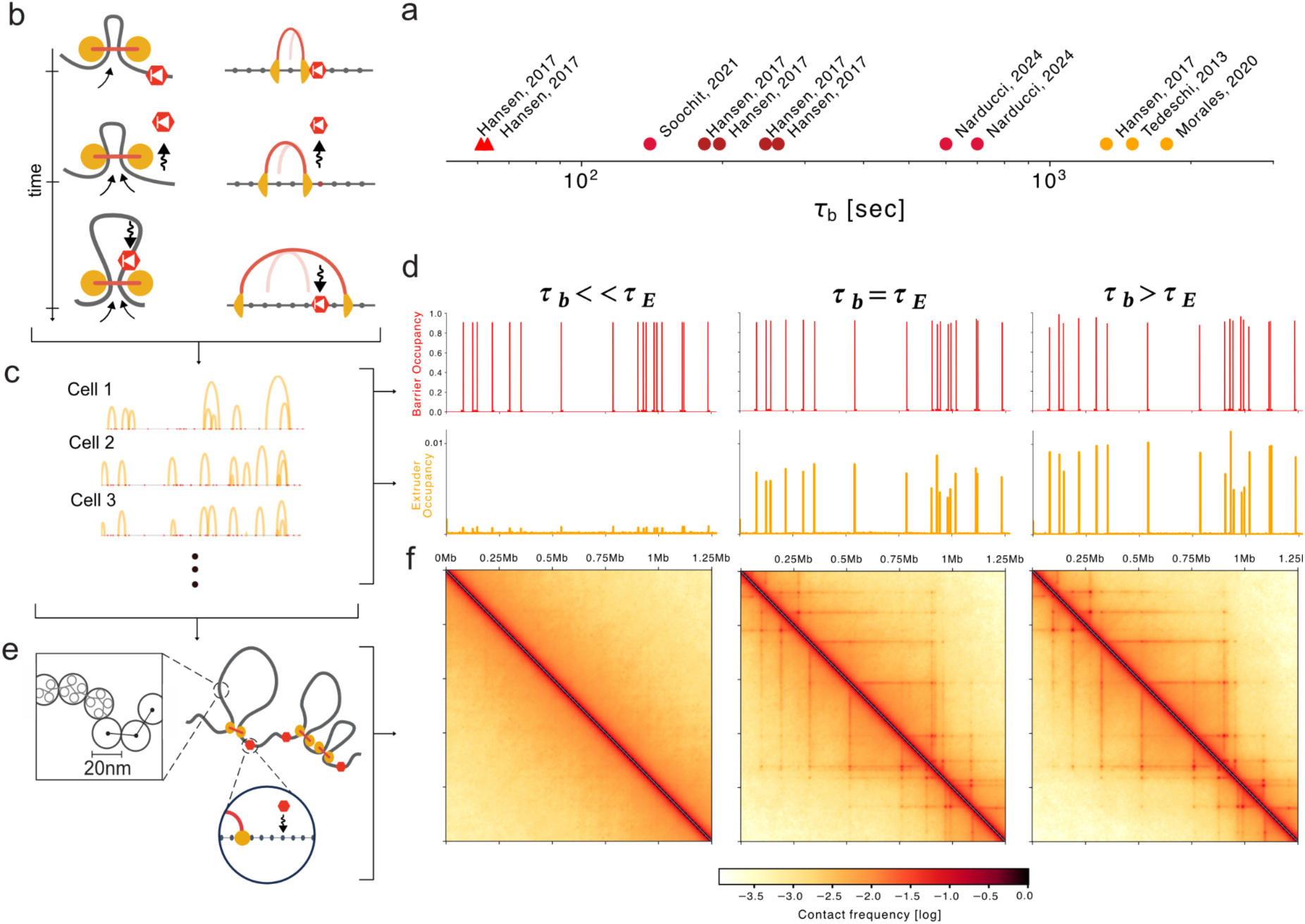
Dynamic barriers show distinct impacts at fixed occupancy. **a.** Residence times for biophysical measurements of CTCF (red) and cohesin (orange) in mouse cells. Measurements for individual cell lines or acquisition rates shown as separate points. Shapes indicate SPT (triangle) or FRAP (circle). See **Table S1** for details. **b.** *Left:* Illustration of CTCF as a dynamic barrier to loop extrusion. *Right:* lattice implementation, where CTCF sites are bound with timescale 𝞃_b_, and unbound with timescale 𝞃_u_. Extruder position at previous time step shown as a light arch. If a barrier becomes unbound (middle), an extruder blocked at this site can continue extruding (bottom). Note CTCF can re-bind when the barrier is inside of an extruded loop. **c.** Snapshots of extruder positions in different cells, where arcs indicate connected left and right legs. To generate *in silico* ChIP-seq tracks, extruder positions are recorded for both legs from 500,000 timepoints, representing a population of cells, and averaged. **d.** *In silico* ChIP-seq for extruders and barriers across a 1.25 Mb region. Barrier bound times 𝞃_b_ (4.1s, 1350s, 4050s) range from much less to much greater than extruder lifetime (𝞃_e_) yet are kept at fixed occupancy (0.9). Extruder ChIP-seq displays strong peaks if 𝞃_b_ >> 𝞃_e_ and very weak peaks if 𝞃_b_ << 𝞃_e_. Note that variation in extruder peak heights comes from differential spacing between barriers, as all barriers have identical parameters in these simulations. **e.** Extruder positions from 1D lattice simulations are used as input to 3D polymer simulations. Ensembles of 3D conformations are generated to produce contact maps. **f.** *In silico* contact maps for the same simulated region and parameters as (**d**) binned to 2.5 kb resolution. Barriers with similar or longer bound times than the extruder lifetime (𝞃_b_ ∼ 𝞃_E,_ 𝞃_b_ >𝞃_E_) display TADs and dots. In contrast, these patterns vanish with transient barriers (𝞃_b_ << 𝞃_E_), even at the same occupancy .

Biophysical simulations enable the simultaneous modeling of protein dynamics and genomic observables *in silico.* Simulations inherently offer independent control over all modeled parameters of the loop extrusion process, including extruder lifetime and barrier bound time. Conformations generated by simulations can be used to extract *in silico* ChIP-seq, Hi-C or imaging. Still, existing models either neglected the dynamics of CTCF barriers (Fudenberg et al. 2016; Gabriele et al. 2022) or were not analyzed in terms of residence times as derived from experimental data (Rossini et al. 2022). Thus, the quantitative impacts of CTCF barrier dynamics on ChIP-seq or Hi-C features remain largely unexplored.

To account for the biophysical kinetics of CTCF, we developed a model for loop extrusion with dynamic barriers that stochastically switch between bound and unbound states. We observed distinct behaviors depending on the ratio of barrier versus extruder lifetimes. Despite the fact that genomic data are population-averaged snapshots, our quantitative comparisons revealed multiple signatures that are consistent with dynamic barriers yet incompatible with static barriers. We additionally found that barrier dynamics modulate observables at all scales of genome organization and from multiple modalities including: (i), the fraction of cohesin at CTCF peaks in ChIP-seq, (ii) the strength of isolation and dots in Hi-C contact maps, to (iii) images of whole-chromosome morphology. Together, our analyses illustrate how biophysically-informed models can be directly confronted with experimental data to sharpen our understanding of genome folding.

## Results

### A model of dynamic CTCF barriers for loop extrusion

To characterize the role of CTCF dynamics for loop extrusion, we extend prior models and explicitly model CTCF occupancy at a set of its cognate sites (**Fig. 1b**). CTCF sites dynamically switch between an occupied bound state (with bound time 𝞃_b_) and unbound state (with unbound time 𝞃_u_). When CTCF sites are occupied, they are unidirectional barriers that prevent loop extruders from passing until they become unbound. We simulate loop extrusion dynamics on a 1D lattice at 250 bp resolution with randomly positioned CTCF sites and cohesin lifetime chosen to approximate experimental estimates in mouse embryonic stem cells (mESC: average barrier distance 75 kb, extruder lifetime 22 minutes; **Methods**). The fine-scale lattice enables modeling CTCF exchanges that are rapid relative to the lifetime of cohesin extruders, while still using the fixed-timestep update scheme used previously (Fudenberg et al. 2016; Nuebler et al. 2018). While in principle each CTCF site could have a different bound and unbound time, to characterize key behaviors of the dynamic barriers model across a broad range of parameters, however, we focused on a scenario where all barriers shared the same dynamic timescales. To simulate chromatin, we modeled 2.5 Mb of DNA as a 50 nm fiber with monomers of size 2.5 kb (**Methods**).

We generated *in silico* ChIP-seq and Hi-C from dynamic barrier simulations. To generate *in silico* ChIP-seq data (**Fig. 1c**) we collected the positions of extruders along the 1D lattice and computed their average frequency at each position. For *in silico* Hi-C maps (**Fig. 1d**), we collected conformations from the 3D simulations and recorded all pairs of monomers with spatial distances below a fixed capture radius as contacts (as previously (Nuebler et al. 2018), **Methods**).

At fixed occupancy, *in-silico* ChIP-seq and Hi-C maps are strongly influenced by the CTCF bound time. At high (90%) occupancy, long-lived barriers result in pronounced peaks in ChIP-seq profiles, as well as clearly-defined TADs and dots in Hi-C maps, reminiscent of experimental observations (**Fig. 1c,d**). In contrast, if the barrier bound time is much smaller than the lifetime of extruders (𝞃_b_ << 𝞃_E_), each of these features largely vanish. To determine the range of barrier bound times in vivo, we developed an analytical understanding of barrier dynamics and then compared simulations with multiple quantitative features of genomic data.

### Analytical approach relating loop sizes to barrier dynamics

To characterize when our simulations would display short versus long barrier bound time behavior, we considered a simplified model that was analytically tractable and quantified how loop sizes depend on barrier dynamics. In the simplified model, a single extruder loads precisely between two convergent barriers. We observed a pronounced decrease of loop size with increasing barrier bound times (**Fig. 2a**). Barrier unbound time played a role as well. For short unbound times, the decreased loop size appeared at a fixed barrier bound time that was slightly smaller than the extruder lifetime. When unbound time exceeded bound time, however, the sharp decrease of loop sizes occurred at progressively longer bound times. Similar behavior was observed for our biologically-informed layout, though loop sizes were lower due to collisions with other extruders as well as sequential encounters with barriers (**Fig. S2a**).

**Figure 2.**
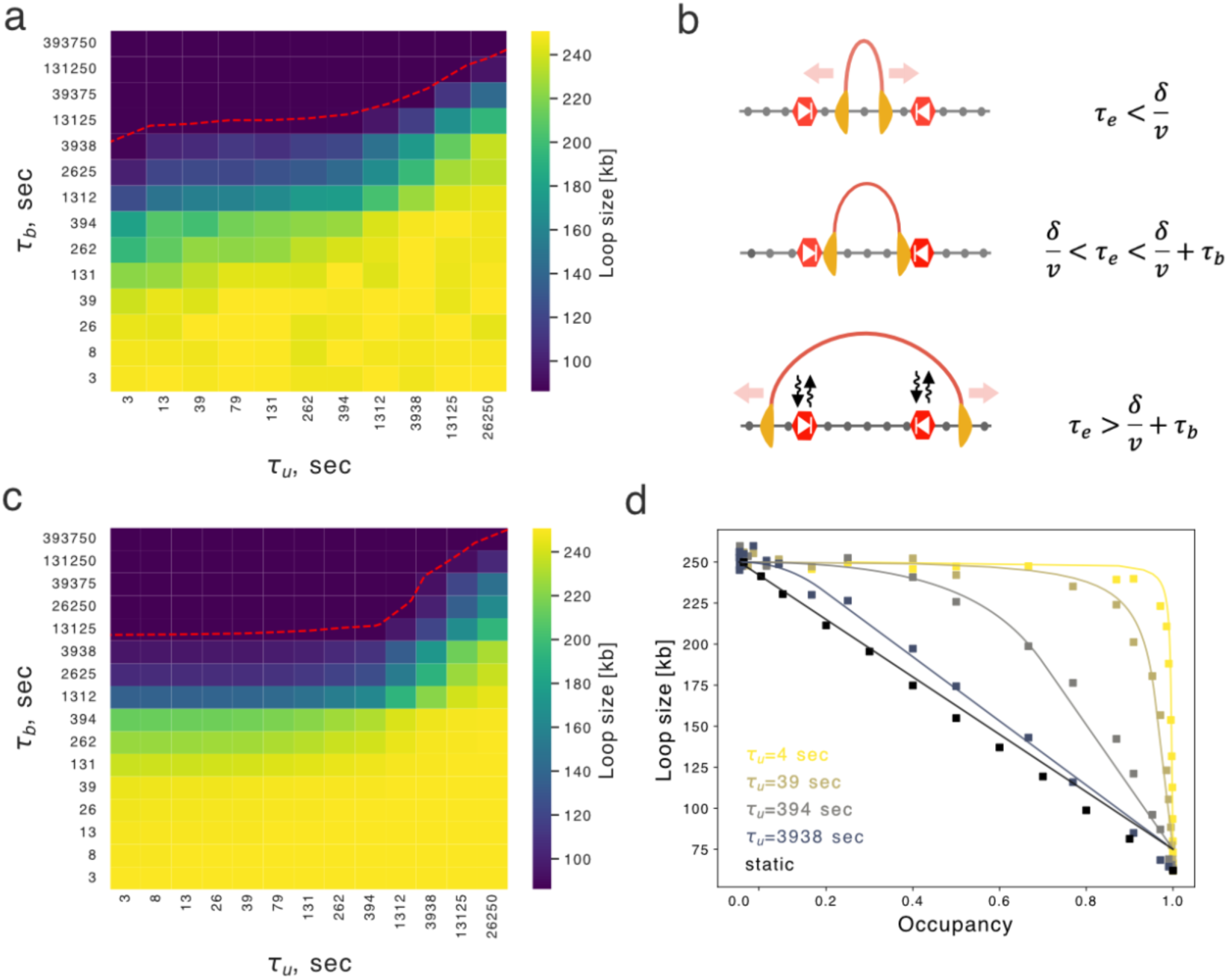
Dynamic barriers lead to distinct regimes for predicted loop size. **a.** Heatmap showing loop sizes for simulations with the simplified layout as a function of bound and unbound times. The simplified layout has an extruder loaded halfway between two convergent barriers. The red dashed contour line indicates loop size = 87.5 kb, equivalent to 𝛿. **b.** Illustration of simplified layout and three regimes of extruder lifetime compared to barrier bound times. Red arrow indicates unimpeded movement of the extruder. At short times (top) the extruder moves unimpeded; at intermediate times (middle) the extruder is stalled; at long times (bottom) the extruder has bypassed the barrier. **c.** As in (a), but computed from the analytical formula. **d.** Loop size versus occupancy for analytical predictions (lines) versus simulations (dots). Note how curves approach the “static” barrier model at large 𝞃_b_ (darker lines), where loop size is a linearly decreasing function of the occupancy.

With this simplified model, three distinct behaviors of the extruder emerge (**Fig. 2b**) depending on extruder lifetime (τ_E_) relative to the barrier distance (δ) divided by extrusion rate (v) and bound time (τ_b_):

1. When extruder lifetime is insufficient to reach the barriers (τ_E_ < δ/v), loop extrusion is unimpeded and loop size is simply proportional to extrusion rate times lifetime.
2. If extruder lifetime is sufficient to reach the barriers but less than the barrier bound time (δ/v < τ_E_ < δ/v + τ_b_), then the extruder bypasses the barrier if it is unoccupied but is blocked if it is occupied.
3. If the lifetime of the extruder exceeds the barrier bound time (δ/v + τ_b_ < τ_E_), the extruder can ultimately bypass the barrier even if its site is occupied. In this case, the barrier unbinds chromatin before the extruder, and we must account for continued loop enlargement after stalling.

Considering extrusion behavior in these three regimes allowed us to derive analytical expressions for loop size as a function of extruder lifetime, bound time, unbound time, barrier separation, and extrusion rate (**Methods**).

These equations capture key aspects of barrier dynamics on extruded loop sizes, as evidenced by the similar heatmap for loop size as a function of bound and unbound time (**Fig. 2a,c**). The quantitative agreement is highlighted in plots of occupancy versus loop size for various unbound times (**Fig. 2d**). Moreover, equations highlight a sharp change in predicted loop size at extruder lifetime minus barrier distance divided by extrusion rate (τ_E_ - 𝛿/v) at short barrier unbound times (**Fig. S2b**). At long unbound times (τ_u_>>τ_E_), the impact of barriers on loop sizes approaches that of static barriers (**Fig. 2d**). Analytical expressions reveal the difference between static barriers, where loop size is only sensitive to occupancy, and dynamic barriers, where the bound time plays a central role in determining loop size. Despite these insights, loop sizes remain challenging to measure directly in experiments.

### CTCF dynamics modulate cohesin positioning along the genome

To characterize the range of CTCF bound times consistent with experimental observations, we considered the predicted impact of dynamic barriers on ChIP-seq, Hi-C, and imaging. Returning to our biologically-inspired layout (**Fig. 1**), we first focused on ChIP-seq due to its high genomic resolution and lower sequencing cost as compared to Hi-C. Simulated ChIP-seq is also less expensive computationally than full 3D polymer simulations, as it is derived from 1D lattice simulations of extruder positions.

Experimentally, the enrichment of cohesin at CTCF sites can be quantified as the fraction of cohesin reads in CTCF peaks (FRiP). We similarly quantified our *in silico* ChIP-seq as the fraction of extruders positioned at barrier locations relative to the total number of extruders (**Fig. 3a**). Higher values indicate more extruder accumulation at barriers.

**Figure 3.**
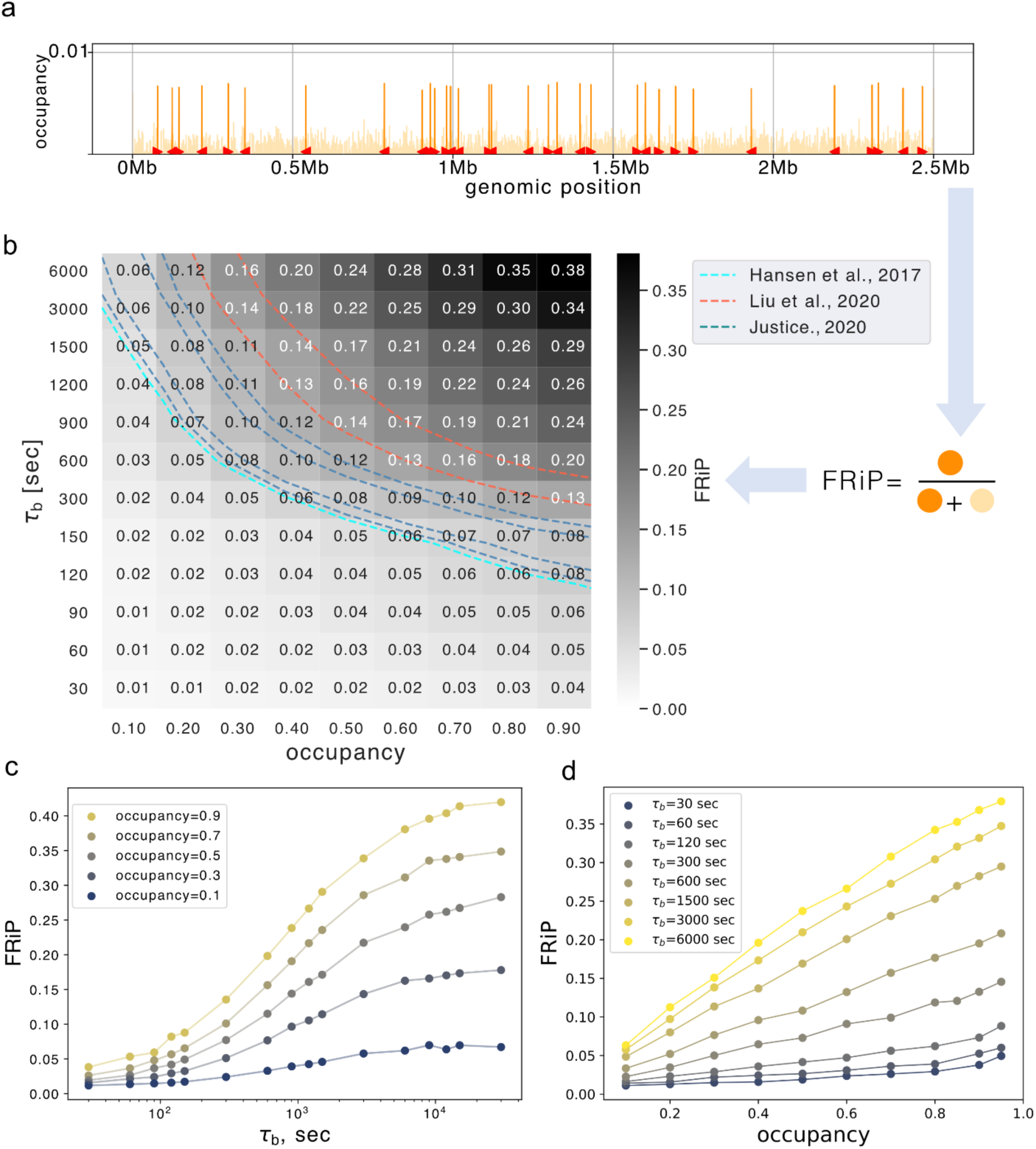
CTCF dynamics modulate cohesin positioning along the genome. **a.** Illustration of how FRiP is calculated for simulations: position of extruder legs are recorded, and then the fraction that overlap CTCF barrier positions (highlighted in orange) is computed. Note that variation in height of individual peaks is due to the differential spacing between peaks, as all barriers are the same strength in these simulations. **b.** Heatmap showing FRiP for a range of CTCF site bound time (𝞃_b_) vs. occupancy. Experimental values for FRiP are depicted with dashed contour lines, ranging from FRiP = 0.07 (Hansen et al. 2017) to 0.16 (N. Q. Liu et al. 2021). **c.** FRiP as a function of 𝞃_b_ for various fixed occupancies. At small 𝞃_b_ FRiP increases rapidly before reaching a plateau. **d.** FRiP as a function of occupancy for various 𝞃_b_. FRiP increases linearly with occupancy at high 𝞃_b_, but remains low at all occupancies when 𝞃_u_ << 𝞃_E_ (=1312 sec).

To characterize the dynamic barrier model, we quantified FRiP across a range of barrier bound time and occupancies (**Fig. 3b, S3a**). We found that FRiP generally displayed the opposite trend as loop size as a function 𝞃_b_: more potent barriers lead to higher FRiP (**Fig. S3b**). At fixed barrier occupancy, FRiP increased with barrier bound time **(Fig. 3c)**. With increasing bound time, FRiP first displays a rapid increase followed by a plateau. This plateau has equivalent FRiP to a model with static barriers (**Fig. S3d**). Still, FRiP increases with occupancy only at high enough barrier bound time and remains negligible when barriers are transient (**Fig. 3d**). As for loop sizes, we observed a transition between high and low FRiP regimes when the barrier bound time approaches the extruder lifetime minus the spacing between barriers (i.e. 𝞃_b_ ∼ 𝞃_E_ - δ_CTCF_/v). The strong dependence of FRiP on barrier bound time in simulations aligns with experimental observations that CTCF lifetime better correlates with CTCF-cohesin peak overlap than the CTCF fraction bound across a set of CTCF zinc-finger mutants (Do et al., 2024).

We next considered how other parameters of our model could modulate FRiP. When we plotted FRiP versus occupancy for various unbound times, we found distinct relationships at short and long unbound times (**Fig. S3c**). At short unbound times, the FRiP versus occupancy curve is concave. At long unbound times, the relationship between FRiP and occupancy is roughly linear. Increasing extruder lifetime increased FRiP for sufficiently high 𝞃_b_ (**Fig. S3e**). In contrast, FRiP only slightly increased at higher extruder separation (**Fig. S3f**). We next tested whether clusters of CTCF sites would yield different FRiP than an isolated site. With a fixed number of bound CTCFs per cluster (i.e., when 𝞃_u_ per barrier was increased proportionally with the number of sites in a cluster), FRiP remained relatively constant (**Fig. S3g,h**).

The strong dependence of FRiP on barrier dynamics in simulations suggested that experimental FRiP could be used to estimate CTCF barrier bound times (see contours for experimental FRiP 0.07-0.15 in **Fig. 3b**). Assuming an average occupancy 0.7 (Sönmezer et al. 2021), the dynamic barriers model argues for a bound time between 150-900 seconds (i.e., 1/10 𝞃_E_ < 𝞃_b_ < 2/3 𝞃_E_). Assuming a slightly lower average occupancy gives a slightly higher estimated bound time. Despite being tractable to compute in simulations, we noticed that experimental estimates for FRiP were quite variable, potentially due to the choice of antibodies for ChIP-seq. We thus considered whether orthogonal genomic datasets could further constrain estimates for barrier dynamics *in vivo*.

### Barrier dynamics modify isolation of neighboring TADs

To determine the impact of dynamic barriers on 3D genome organization, we quantified patterns in simulated contact maps across a range of barrier bound times. We first quantified the isolation imposed by dynamic barriers on upstream and downstream regions. We defined isolation as the relative frequency of contacts in triangular regions within TADs versus between neighboring TADs (**Fig. 4a, Methods**). We found a strong dependence of TAD isolation on CTCF dynamics (**Fig. 4b**). As for FRiP, isolation varied with both barrier bound times and unbound time, even at fixed occupancy rates. Importantly, high occupancy alone was insufficient to produce strong TADs in simulated Hi-C maps, as sufficient barrier bound times are also required. Comparison with an average experimental isolation score and occupancy suggests barrier bound times approaching that of the extruder lifetime (900-1500 seconds).

**Figure 4.**
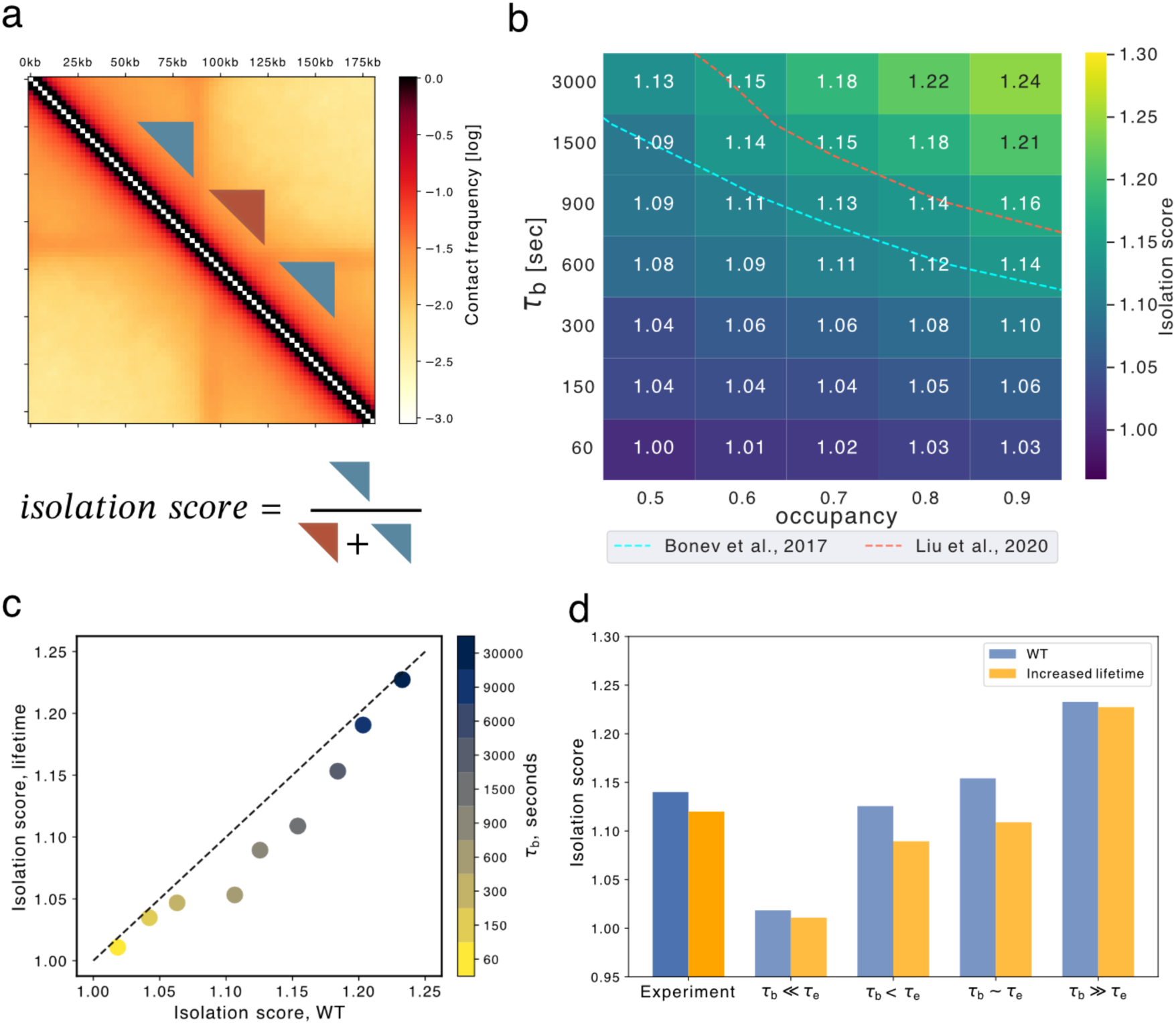
Dynamic barriers modify isolation of neighboring TADs, even at a fixed occupancy rate. **a.** Illustration of how the isolation score is calculated using a window size of 50 kb, and equal size triangular areas within (blue) and between (brown) TADs. **b**. heatmap showing sweeps of isolation score for barrier bound time (𝞃_b_) versus occupancy, averaged across all simulated barriers. At large 𝞃_b_ and high occupancy, TADs are more isolated. Experimental isolation scores shown as dashed lines (1.12 and 1.14, also see **Figure S4**) **c.** Comparison of isolation scores between normal and threefold increased extruder lifetime conditions for various barrier bound time at occupancy 0.7, with point size proportional to the bound time. **d.** Comparison between isolation scores for experiments before (blue) and after WAPL depletion (yellow), along with simulations with WT (blue) and higher (yellow) lifetime for various 𝞃**_b_**.

Intriguingly, when we performed simulations with increased extruder lifetimes, we observed lower isolation between TADs for intermediate barrier bound times (**Fig. 4c,d**). Similar behavior was observed in experimental Hi-C for WT versus WAPL depletion conditions (**Fig. 4d, Fig. S4c**). Hi-C data thus displays a signature of dynamic barriers, as isolation score does not decrease in the static barrier regime (i.e., 𝞃_b_ >> 𝞃_E_, **Fig. 4c,d**). The impact of extruder lifetime on isolation contrasts with its effect on FRiP (**Fig. S4d**). While FRiP and isolation score are correlated for fixed extruder parameters, they are anti-correlated across differing extruder lifetimes, presenting an instance of Simpson’s paradox (Blyth 1972). We hypothesize this arises because isolation score depends on all polymer positions between barriers, whereas ChIP-seq FRiP depends primarily on encounters directly at the barriers.

### Hi-C dot strength versus genomic distance requires dynamic boundaries

We next investigated how the strength of Hi-C dots depended on simulated CTCF barrier dynamics. As dots correspond to pairs of positions with accumulated extruders, we hypothesized that dot strength might be more closely correlated with FRiP than TAD isolation. We defined dot strength as the frequency of contacts between pairs of convergent barriers relative to their surrounding areas (**Fig. 5a, Methods**, as previously (Rao et al. 2014; Flyamer, Illingworth, and Bickmore 2020) and computed dot strengths across a range of barrier parameters (**Fig. 5b**). Comparison with experimental data suggested a barrier bound time of 900-1500 seconds, similar to the estimate from isolation scores. Similarly to both FRiP and TAD isolation, we observed that dot strength depended strongly on barrier bound time, even at constant occupancy (**Fig. 5b**). Despite being calculated from the same simulated Hi-C maps, we observed a different behavior for dot strength versus isolation scores as a function of extruder lifetime. Indeed, for higher extruder lifetimes we found that dot strength was increased at all barrier bound times (**Fig.** 5c-d, S5a-b).

**Figure 5.**
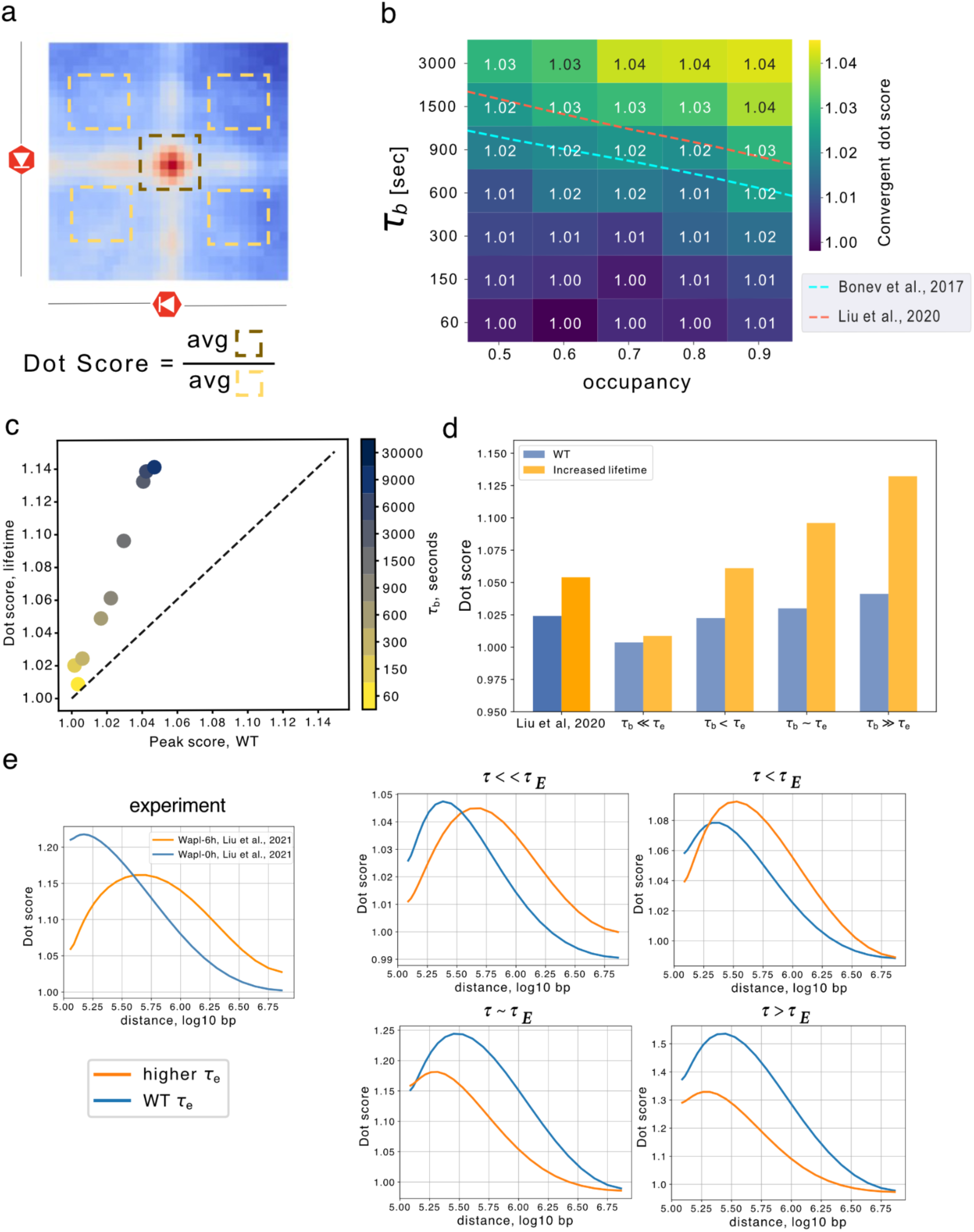
Hi-C dot strength versus distance requires dynamic boundaries. **a.** Dot scores were calculated between pairs of convergent barriers. First, 80 kb snippets centered around barrier pairs were collected from maps binned at 10 kb resolution. Genomic distances were binned into 25 logarithmically spaced bins between 100 kb to 7500 kb, and snippets in each bin were averaged. Dots scores were computed as the ratio of contacts in the snippet center relative to four control regions. **b.** Heatmap of averaged dot score for convergent pairs of barriers over all distances. Note higher scores for larger 𝞃_b_ at fixed occupancies. Dashed lines indicate experimental dot scores (blue 1.0214, red 1.0233). **c.** Averaged convergent dot score for WT extruders versus higher lifetime extruders, indicating stronger dots for the latter, particularly at higher 𝞃_b_. **d.** Comparison between convergent dot scores for experiments before (blue) and after WAPL depletion (yellow), along with simulations for various 𝞃**_b_** with reference (blue) or increased (yellow) lifetime. Experimental dot scores increase after WAPL depletion (from 1.0233 to 1.0492). **e.** Convergent dot score as a function of distance, for extruders with either WT (blue) and higher (orange) lifetimes at three CTCF bound times. Only simulations with 𝞃_b_<𝞃_E_ displayed similar behavior to experiments, where the higher-lifetime curve starts lower and has a peak after the wildtype curve.

A distinct aspect of dot strength from FRiP or isolation is that it can be computed as a function of the genomic distance between dot anchors. In simulations with high barrier bound times, increased extruder lifetime enhanced average dot strength across all genomic distances. However, experimentally increasing extruder lifetime via WAPL depletion does not substantially increase the maximal average dot strength (**Fig. 5d, Fig. S5d**). Instead, the maximal dot strength in dWAPL shifts to further genomic distances, with weaker dot strengths than WT condition at moderate (100 kb) genomic distances. To characterize this behavior in simulations, we generated contact maps for a layout representing 10-times more chromatin to provide a sufficient number of dot anchors for analysis at short genomic distances. In simulations, we found the shift to further genomic distances for higher lifetime extruders can be observed when barriers display more rapid dynamics (𝞃_b_= 150-300 seconds, **Fig. 5e**), yet cannot be observed in models with static barriers (**Fig. S5e**). Dynamic barriers also better reproduced the experimentally observed magnitude of the dot strength versus distance curves than static or very long-lived barriers (**Fig. S5c)**. Dot strength versus distance thus constitutes a new Hi-C signature of barrier dynamics *in vivo*.

### Local barrier dynamics can shape whole-chromosome morphology

When cohesin lifetime on chromatin is experimentally increased following WAPL depletion, chromosomes condense and assume a vermicelli-like morphology (Tedeschi et al. 2013). The axis of vermicelli chromatids are enriched for both cohesin and CTCF. While a similar vermicelli morphology can emerge with increased extruder lifetime in simulations,, the impact of barrier dynamics on vermicelli formation have not been characterized. If we increased extruder lifetime 7-fold (as estimated experimentally (Wutz et al. 2017)) with transient barriers (𝞃_b_ = 30 seconds, occupancy 0.7), we observed vermicelli formation in simulations (**Fig. 6a**). At that same extruder lifetime and barrier occupancy, if we increased barrier bound time to that approaching the static regime (𝞃_b_= 4500 seconds), we no longer observed vermicelli formation. To quantify vermicelli formation, we computed the Pearson correlation between the spatial positions of extruders and chromatin in 3D (**Methods,** Tortora and Fudenberg 2024). We then assayed simulated vermicelli formation across a range of extruder lifetimes and barrier bound times for fixed barrier occupancy. At WT extruder lifetimes, vermicelli are not present and there is a low spatial overlap between cohesin and chromatin. At higher extruder lifetimes, a vermicelli morphology is observed. However, if barrier bound times are also high, vermicelli formation is impeded. The experimental observation of vermicelli after an estimated 7-fold increase in lifetime therefore allows us to use simulations to estimate an upper bound on the barrier bound time of 600 seconds (∼½𝞃_E_). As for FRiP, isolation, and dot strength, vermicelli formation depended on barrier bound time at fixed occupancies. How dynamic barriers enable vermicelli formation can be understood by considering the case where two CTCF sites are close together without an extruder in the interval: instead of relying on a low probability event that an extruder is loaded between the barriers, this gap can be closed if an extruder is released after either barrier unbinds (**Fig. 6b**). In sum, vermicelli formation relies on a delicate balance between barrier dynamics and the extruder lifetime.

**Figure 6:**
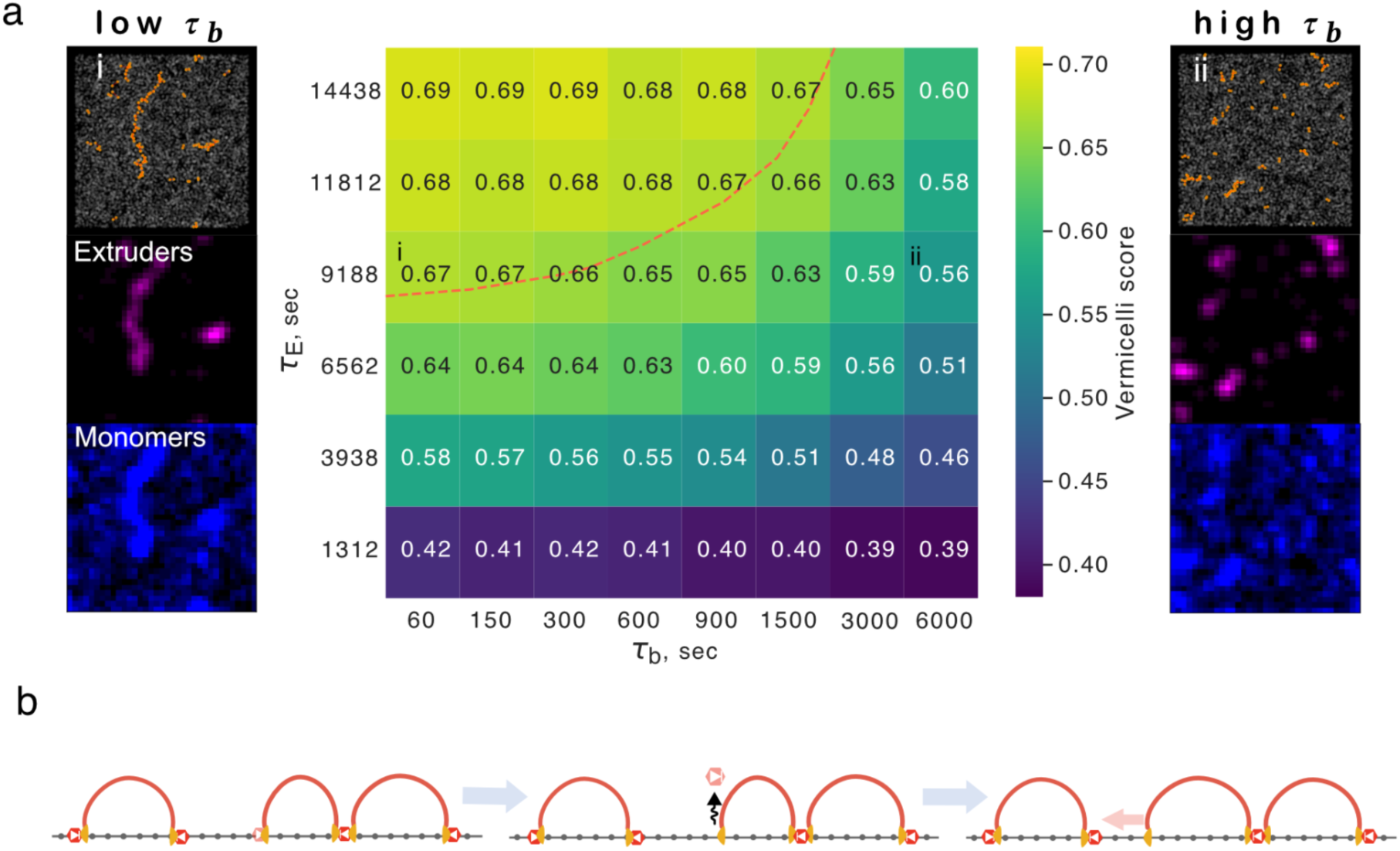
Local barrier dynamics can shape whole-chromosome morphology. **a.** Heatmap indicating vermicelli conformation at various barrier bound times 𝞃_b_ relative to extruder lifetime (𝞃_E_) at fixed barrier occupancy (0.7). The red dashed line indicates parameter sets displaying prominent vermicelli (scores above 0.65). *Left column:* representative conformation above simulated microscopy for chromatin (blue) and extruders (purple) at 7-fold increased extruder lifetime approximating WAPL depletion and very short barrier bound time (𝞃_b_<< 𝞃_E_), position in heatmap indicated by (i). *Right column:* similarly, albeit for very long barrier bound time (𝞃_b_>>𝞃_E_), with position in heatmap indicated by (ii). **b.** Illustration of how dynamic barriers can have global consequences for vermicelli formation by enabling gap closure between consecutive loops. In this three-step process, an extruder is halted at a barrier next to a gap; the barrier unbinds, allowing extruder bypass; finally, the gap can be closed, irrespective of whether the barrier re-binds.

## Discussion

By characterizing a dynamic CTCF barrier model for interphase loop extrusion, we find that barriers with identical occupancy can yield distinct predictions for ChIP-seq, TADs, corner dots, and chromosome morphology. Model behavior can be summarized as having three regimes depending on the ratio of CTCF binding time (𝞃_b_) to extruder lifetime ( 𝞃_E_):

i. *Transient barriers* (𝞃_b_ << 𝞃_E_): extruders easily bypass barriers.
ii. *Dynamic barriers* (𝞃_b_ ⪅ 𝞃_E_): extruders stall at barriers with occasional release.
iii. *Quasi-static barriers* (𝞃_b_ >> 𝞃_E_): when a barrier is present, it almost permanently stalls extruders. Only the dynamic barriers regime, with barrier bound times of 10-15 min (i.e. 𝞃_b_ ∼ ½ 𝞃_E_ to ⅔ 𝞃_E_), produced ChIP-seq FRiP, TADs, dots, and vermicelli in good agreement with experimental data.

The dynamic barriers model predicts that barrier bound time is more important than occupancy for determining many features observed in experiments. Indeed, barriers required sufficient bound time to reliably block and accumulate extruders. This insight from simulations could provide an alternative explanation for why many CTCF ChIP-Seq peaks (20-50%) do not display corresponding RAD21 peaks, beyond potential issues with antibody quality (Pugacheva et al. 2020). Moreover, the dynamic barriers model predicted a stronger relationship between cohesin accumulation and barrier bound time than barrier occupancy. A similar trend was recently observed experimentally for a panel of CTCF zinc-finger mutants (Do et al. 2024).

Dynamic barriers are additionally required to explain three features of genome folding after WAPL depletion, where the lifetime of cohesin loop extruders is thought to greatly increase (Wutz et al. 2017; Tedeschi et al. 2013; Haarhuis et al. 2017). First, in dWAPL cells the maximal dot strength shifts to further genomic distances. This can be recapitulated with dynamic barriers, yet not with static barriers. Second, WT Hi-C maps indicate stronger isolation compared to dWAPL, which again cannot be recapitulated by static barriers. Third, dWAPL chromosomes adopt a vermicelli morphology. Dynamic barriers allow bypass and vermicelli formation, yet static barriers disrupt vermicelli formation even by very high lifetime extruders. An intriguing consequence of dynamic barriers is that longer-lived extruders could potentially bypass multiple barriers sequentially, equalizing contact frequencies across a large genomic region (**Fig. 7**). We hypothesize that this feature of dynamic barriers is harnessed to diversify promoter choice at the protocadherin locus and specify neuronal identity, as recently reported (Kiefer et al. 2023).

**Figure 7.**
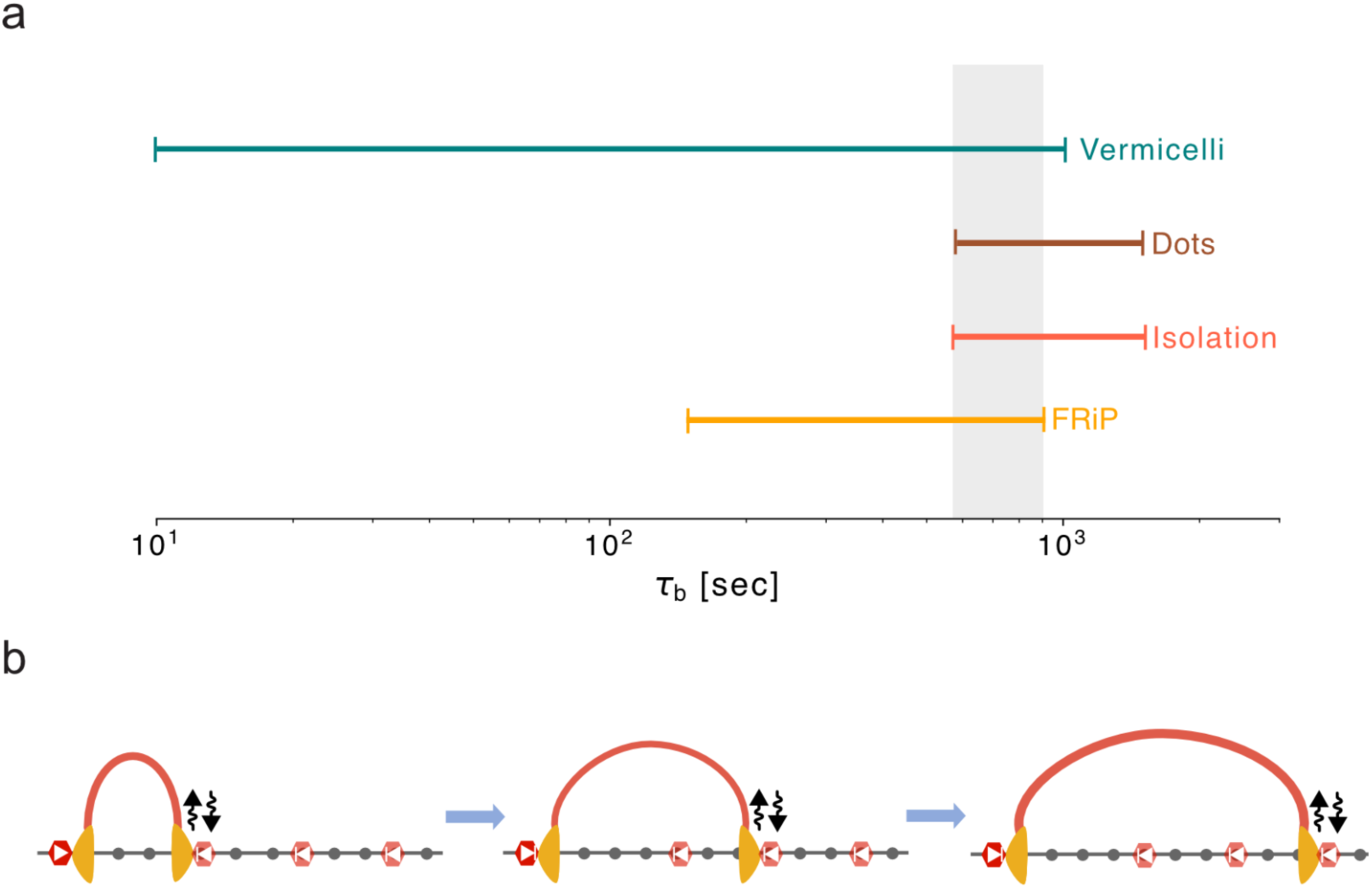
Dynamic barriers are required for agreement with multiple features of experimental data. **a.** Range of barrier bound times from the dynamic barriers model in agreement with experimental data based on FRiP, isolation score, dot score, and vermicelli formation. Highlighted area (grey) indicates region of agreement across metrics (𝞃_b_ between 10-15 minutes). **b.** Consequence of dynamic barrier model: long-lived extruders can sequentially bypass a series of dynamic barriers. This can equalize contact frequencies between a long-lived barrier, acting as an anchor, (left site, dark red) and multiple downstream dynamic barrier positions (right three sites, lighter red).

While FRAP and SPT are insightful for their ability to measure CTCF bound times in vivo, both techniques extract average times across the genome. Across the genome, however, individual sites appear to have a range of biophysical properties (Luan et al. 2021). In addition to site-specific bound times and occupancy (Sönmezer et al. 2021; Ramani, Qiu, and Shendure 2019), CTCF-RNA associations (Hansen et al. 2020) could provide site-specific unbound times as well.

Despite considering a simplified scenario, where all sites had the same bound and unbound times, our estimated bound times are only slightly larger than those reported via the latest FRAP measurements (∼12 min versus ∼10 min (Narducci and Hansen 2024)). If we assumed a slightly lower occupancy (50% rather than 70%), our estimated bound times would increase (to ∼25 min). Incorporating additional details into future models of barrier dynamics could thus improve agreement with experimental measurements. First, site-specific CTCF bound times could enable a subpopulation of longer-lived CTCFs to provide most of the barrier functionality, yet be minimally reflected by current SPT or FRAP assays.

Support for this comes from reports of stably-bound CTCF populations (52.7 minutes in resting and 27.1 minutes in activated B-cells (Kieffer-Kwon et al. 2017), 16.7 minutes in a human fibroblast cell line (Agarwal et al. 2017)) as well as persistent protection of CTCF motifs after long MNase digestions (20min, (Ramani, Qiu, and Shendure 2019)). Second, CTCF barrier activity involves a more complex mechanism than simply blocking extruders (Hansen 2020), perhaps via protection from WAPL (Li et al. 2020; Y. Liu and Dekker 2022; Wutz et al. 2020; Brunner et al. 2024), preventing association with NIPBL (Rhodes et al. 2017), interaction with PDS5 (Wutz et al. 2017; Nora et al. 2020), or via a tension-dependent mechanism (Davidson et al. 2023). Indeed, if CTCF can deactivate the cohesin motor (Hansen 2020), its effective barrier bound time would be longer than that inferred from its dynamics alone.

Given the central role of barrier bound time we reveal, further progress in modeling interphase genome folding will benefit from new experimental approaches to measure site-specific residence times, perhaps by adapting methodology from emerging microscopy approaches (Pomp, Meeussen, and Lenstra 2024). Our work highlights the ability of models to triangulate across multiple experimental modalities, revealing signatures of dynamic barriers evident even in the fixed-time snapshots provided by genomic datasets. In sum, understanding the balance between barrier and extruder dynamics will sharpen our understanding of how cells harness loop extrusion to achieve myriad outcomes.

## Methods

Our simulations of loop extrusion with dynamic barriers integrated a 1D lattice model for loop extrusion with 3D polymer simulations of chromatin. Together this requires specifying the following timescales:

- CTCF site bound time: 𝞃_b_
- CTCF site unbound time: 𝞃_u_
- Extruder lifetime: 𝞃_Extruder_
- Extruder step rate: Δ𝞃_Extruder_
- Lattice update timescale: 𝛿𝑡_*lattice*_
- Polymer update timescale: 𝛿𝑡*_3d_*

### 1D lattice model of loop extrusion

We simulated a one-dimensional lattice with 10,000 sites, each representing 250 bp. At each time step, extruders that were not stalled at barriers took outward steps with 10% probability. Together this implementation enabled fixed-timestep simulations with rapid CTCF exchange (CTCF bound time up to ∼20-30x lower than cohesin), at close to nucleosome resolution. Loop extruders are loaded randomly on the lattice, with a number depending on the separation parameter. For our simulated WT condition this was 250kb (1000 sites), close to that in previous simulations (240kb, (Gabriele et al., 2022)). A loop extruder occupies two lattice sites, one for its right and one for its left leg. Loop enlargement is modeled by re-assigning legs to new lattice sites, where an extruder at (i, j) gets reassigned to (i-1, j+1). Legs of different cohesins cannot bypass each other when they meet. Extruders also cannot exit at either the first or last lattice site. We considered purely unidirectional barriers: only cohesins whose extrusion direction clashes with the barrier orientation can be stalled. If a leg is stalled at an occupied barrier, it no longer moves until that barrier unbinds the lattice, while the other leg can still move independently. Eventually, extruders dissociate from the lattice with a fixed rate determined by the extruder lifetime, which also governs processivity (the average extruded distance without collision), and are loaded randomly at a new position.

To speed up averaging, we used 10 consecutive replicas of the same layout of 10000 sites, giving a full lattice of 100000 sites. Each replica had the same positions of 32 randomly-generated barrier positions, with 16 right-stalling barriers at: [314, 579, 1195, 3717, 3772, 3921, 4451, 5193, 5723, 6302, 6574, 6779, 7000, 9232, 9310, 9861], and 16 left-stalling barriers at: [495, 865, 1404, 2164, 3143, 3615, 3971, 4069, 4480, 4938, 5300, 5587, 6401, 7725, 8764, 9619]. On average, this corresponds to 75 kb (300 sites) between barriers, approximating experimental estimates (74.1kb from ∼217,000 CTCFs per nucleus, 50% bound, and the average genomic content of ∼3*2.7 Gb (Cattoglio et al. 2019)).

We performed 10,000 update steps to generate arrays of extruder positions over time. In wild-type (WT), or reference, conditions extruders had a processivity of 1000 lattice steps (250 kb) and separation of 1000 sites. We initialized simulations with barrier occupancy states chosen proportional to their barrier binding time relative to the sum of their binding and unbinding time.

### 3D polymer simulations

We modeled 2.5 Mb of chromatin as a 50 nm fiber, where each monomer represents 2.5 kb (i.e., 10 1D lattice sites). We chose this resolution for the 3D simulations to reduce computational cost yet still be below a typical experimental Hi-C analysis resolution of 10kb. We used *polychrom* (https://github.com/open2c/polychrom) to implement polymer models, which harnesses OpenMM (Eastman et al. 2017) to generate large ensembles of conformations. We introduced an extra harmonic bond between the two monomers connected by each pair of extruder arms (i, j). When the loop extrusion factor (LEF) advances, this bond is removed and replaced by a new bond between (i-1, j+1). We observe Rouse dynamics for our wildtype scenario and calibrated simulation timescales with experimental timescales by aligning Mean Square Displacement (MSD) curves. Assuming a diffusion coefficient 𝐷 = *0*.*01* 𝜇𝑚*^2^*/𝑠*^0^*^.*5*^ (as in (Nuebler et al. 2018)) with the monomer size used in these simulations (50 nm) each simulated time step in 3D represents 𝛿𝑡*_3_*_’_ = 0.015 sec. We employed 175 3D steps between each 1D lattice update, so a lattice timestep 𝛿𝑡_*lattice*_=17.5 𝛿𝑡*_3d_* amounted to about 2.6 sec of physical time, yielding a stepping rate of 2*250bp / 2.6sec = 190bp/sec. Thus, the WT extruder processivity of 500 lattice steps for each leg translated to an extruder lifetime of 21.8 min, consistent with experimental measurements for cohesin (Hansen et al., 2017). The timescales of barriers (binding and unbinding times, 𝞃_b_ and 𝞃_u_ ) are then calibrated compared to the extruder lifetime. Other parameters in our 3D simulations, including pairwise interaction potential, persistence length, and Langevin integration rate, were the same as those in (Nuebler et al. 2018)

### Analytical formula for loop size

Considering extrusion behavior in for the three regimes described in the main text result in the following equations for the loop size

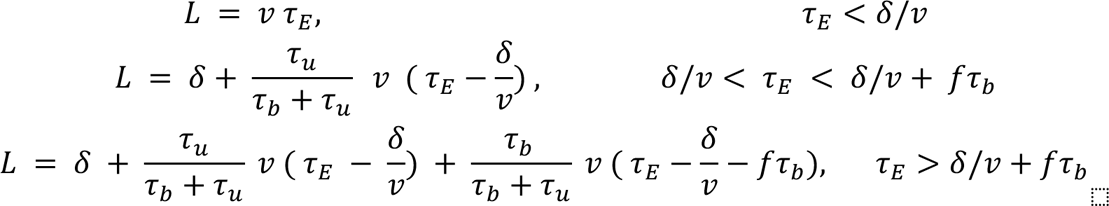

In terms of extruder lifetime (τ_E_), extrusion rate (v), barrier bound time (τ_b_), unbound bound time (τ_u_), and the distance between barriers (𝛿). As a barrier may be bound before the arrival of the extruder, we account for this as the fraction of the bound time, *f* (0<= f <=1), that had elapsed prior to the arrival of the extruder. Loop size can then be obtained by integrating over *f*:

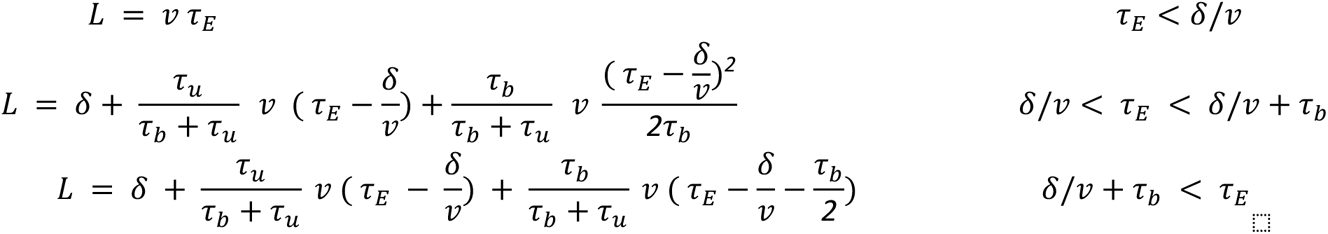

### ChIP-seq and FRiP analysis

We computed in silico ChIP-seq by collecting the positions of extruder legs from a total of 10,000 instances of the 1D lattice simulations. We tabulated extruder leg counts per lattice site, averaged over all ten replicas. From in silico ChIP-seq, we then computed the Fraction of Reads in Peaks (FRiP) as the sum of extruder legs at barrier positions, relative to the total number of extruder legs.

To quantify experimental ChIP-seq data, we analogously calculated FRiP for cohesin ChIP-seq reads in CTCF peaks, using the *fastaFRiP* pipeline (https://github.com/Fudenberg-Research-Group/fastaFRiP), deployed with *Snakemake* (Mölder et al. 2021). Briefly, we aligned FASTQ data to the mm10 genome with *bowtie2* (Langmead and Salzberg 2012) and called peaks with *MACS2* (Zhang et al. 2008). Using the default narrowPeak width for CTCF peaks, we then counted the number of cohesin reads in CTCF peaks with *deeptools* (Ramírez et al. 2016), and divided by the total number of mapped reads to obtain experimental FRiP per dataset.

### Hi-C analysis

For *in silico* Hi-C, we generated a total of 10,000 conformations from polymer simulations. From these conformations, we created Hi-C maps at a resolution of one monomer size (2.5 kb) using the default *polychrom* capture radius (2.3 monomers) to record pairs of monomers in contact, approximating literature values (McCord, Kaplan, and Giorgetti 2020). To improve statistical accuracy, we calculated contacts within subchains of 3,000 monomers for conformations after the system reached a steady state, and averaged the results across subchains and replicates. For a consistent analysis between simulated and experimental data, we converted simulated contact maps to *cooler* (Abdennur and Mirny 2020) format using *polykit* (https://github.com/open2c/polykit). Quantitative analysis of contact maps, including computation of dot and isolation scores, was performed using *chromoscores* (https://github.com/Fudenberg-Research-Group/chromoscores/), which first extracts snippets from simulated maps around barriers or pairs of barriers. Isolation and dot scores are then computed on average snippets from maps binned to 10kb resolution. To compute the isolation score from an average snippet centered on the barrier, we used triangular regions (window size 50kb) upstream, downstream, and spanning the barrier. This choice weights contacts at each distance similarly for the within and between-domain areas, and results in a consistent score for either observed or observed/expected maps. To compute dot scores, we computed average snippets (window size 80 kb) centered around pairs of barriers as a function of genomic distance in 25 logarithmically spaced bins between 100 kb to 7500 kb. For convergent dot scores, we only used snippets around pairs of convergently oriented barriers. From the average snippets, dot scores as a function of distance were computed as the ratio of contacts in the center relative to four control regions. To obtain a single dot score, we computed the average across all genomic distances, weighted by the number of snippets at each genomic distance to account for variation in the number of snippets as a function of genomic distance.

We analyzed experimental Hi-C at the same 10 kb resolution as simulated data. We used Open2C tools *cooler* (Abdennur and Mirny 2020), *pairtools* (Open2C et al. 2023) and *distiller* (Goloborodko et al. 2022) to process reads into binned Hi-C contact maps. To define experimental CTCF barrier positions for isolation and dot analysis, we obtained the overlap between positions of CTCF motifs based on their score from JASPAR (Rauluseviciute et al. 2024) and ChIP-seq peaks (from (Justice et al. 2020)). We then considered the strongest CTCF motif in any 10kb bin (based on their score from *MACS2*), resulting in 29986 positions across the genome. We computed dot and isolation scores from experimental Hi-C data entirely analogously to simulations by using *coolpuppy* to extract snippets and create pile-ups (Flyamer, Illingworth, and Bickmore 2020).

### Vermicelli analysis

The Pearson “vermicelli score” quantifies the spatial relationship between DNA and cohesin positions within a 3D space. We performed a 3D rasterization of these positions into a voxel grid and subsequently applied Gaussian smoothing to the grids. The "vermicelli score" was then calculated by computing the Pearson correlation between the smoothed voxel grids of monomers and extruder positions.

### Code

Code specifying simulations and for performing analysis are available at: https://github.com/Fudenberg-Research-Group/dynamic_extrusion_boundaries.

## Acknowledgements

The authors thank Anders Hansen and Erika Anderson for detailed feedback, and members of the Fudenberg group for helpful discussions. The authors are supported by NIGMS R35GM143116 to GF.

