## Supplemental Information for "Dynamic barriers modulate cohesin positioning and genome folding at fixed occupancy"

### Supplementary Information for: Dynamic barriers modulate cohesin positioning and genome folding at fixed occupancy

Hadi Rahmaninejad[1], Yao Xiao[1], Maxime M.C. Tortora[1], Geoffrey Fudenberg[1]

| paper | protein | condition | cell-type/cell-line | method | bound time, s |
|---|---|---|---|---|---|
| Hansen, 2017 | CTCF | WT | mESC C59 | SPT | 61 |
| Hansen, 2017 | CTCF | WT | mESC C87 | SPT | 63 |
| Hansen, 2017 | CTCF | WT | mESC, C87 | FRAP (11min 0.5hz) | 247 |
| Hansen, 2017 | CTCF | WT | mESC, C87 | FRAP (5min, 1hz) | 183 |
| Hansen, 2017 | CTCF | WT | mESC, C59 | FRAP (11min 0.5hz) | 263 |
| Hansen, 2017 | CTCF | WT | mESC, C59 | FRAP (5min, 1hz) | 197 |
| Hansen, 2020 | CTCF | CTCF-RBRi | mESCC59 | FRAP (11min 0.5hz) | 220 |
| Soochit, 2021 | CTCF | WT | mESC | FRAP (5min, 0.2hz) | 140 |
| Soochit, 2021 | CTCF | CTCF-del8 | mESC | FRAP (5min, 0.2hz) | 17 |
| Soochit, 2021 | CTCF | CTCF-del9 | mESC | FRAP (5min, 0.2hz) | similar to WT |
| Soochit, 2021 | CTCF | CTCF-del10 | mESC | FRAP (5min, 0.2hz) | similar to WT |
| Soochit, 2021 | CTCF | CTCF-del11 | mESC | FRAP (5min, 0.2hz) | similar to WT |
| Narducci, 2024 | CTCF | WT | mESC, clone A | FRAP (10.5min, 0.25hz) | 700 |
| Narducci, 2024 | CTCF | WT | mESC, clone B | FRAP (10.5min, 0.25hz) | 700 |
| Narducci, 2024 | CTCF | WT | mESC, clone D | FRAP (10.5min, 0.25hz) | 600 |
| Narducci, 2024 | CTCF | WT | mESC, C87 | FRAP (10.5min, 0.25hz) | 700 |
| Tedeschi, 2013 | RAD21 | WT | MEF | FRAP | 1500 |
| Hansen, 2017 | RAD21 | WT | mESC | FRAP | 1320 |
| Morales, 2020 | RAD21 | WT | MEFs | FRAP | 1776 |

**Table S1. Experimentally reported SPT and FRAP for CTCF and RAD21 in mouse cells.**

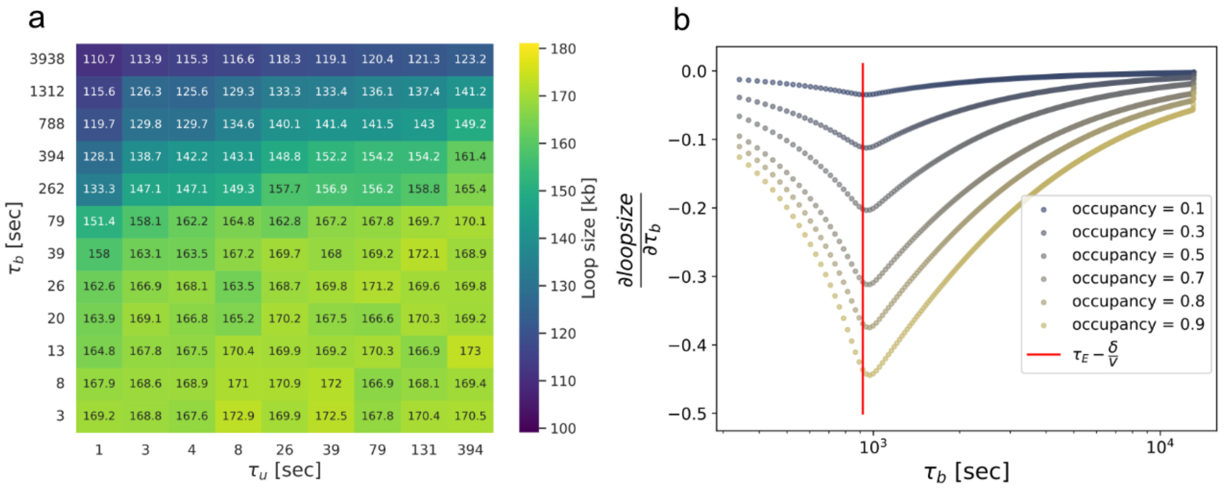


**Figure S2. Loop size dependency on extruder parameters.**

**a.** Heatmap of averaged loop size for simulations with multiple extruders and barriers, the main configuration used throughout the paper. This indicates similar behavior in loop size dependency on binding time, albeit lowered by collisions.

**b**. Derivative of loop size as a function of $\boldsymbol{\tau}_b$, showing a minima at $\boldsymbol{\tau}_E$-δ/v.

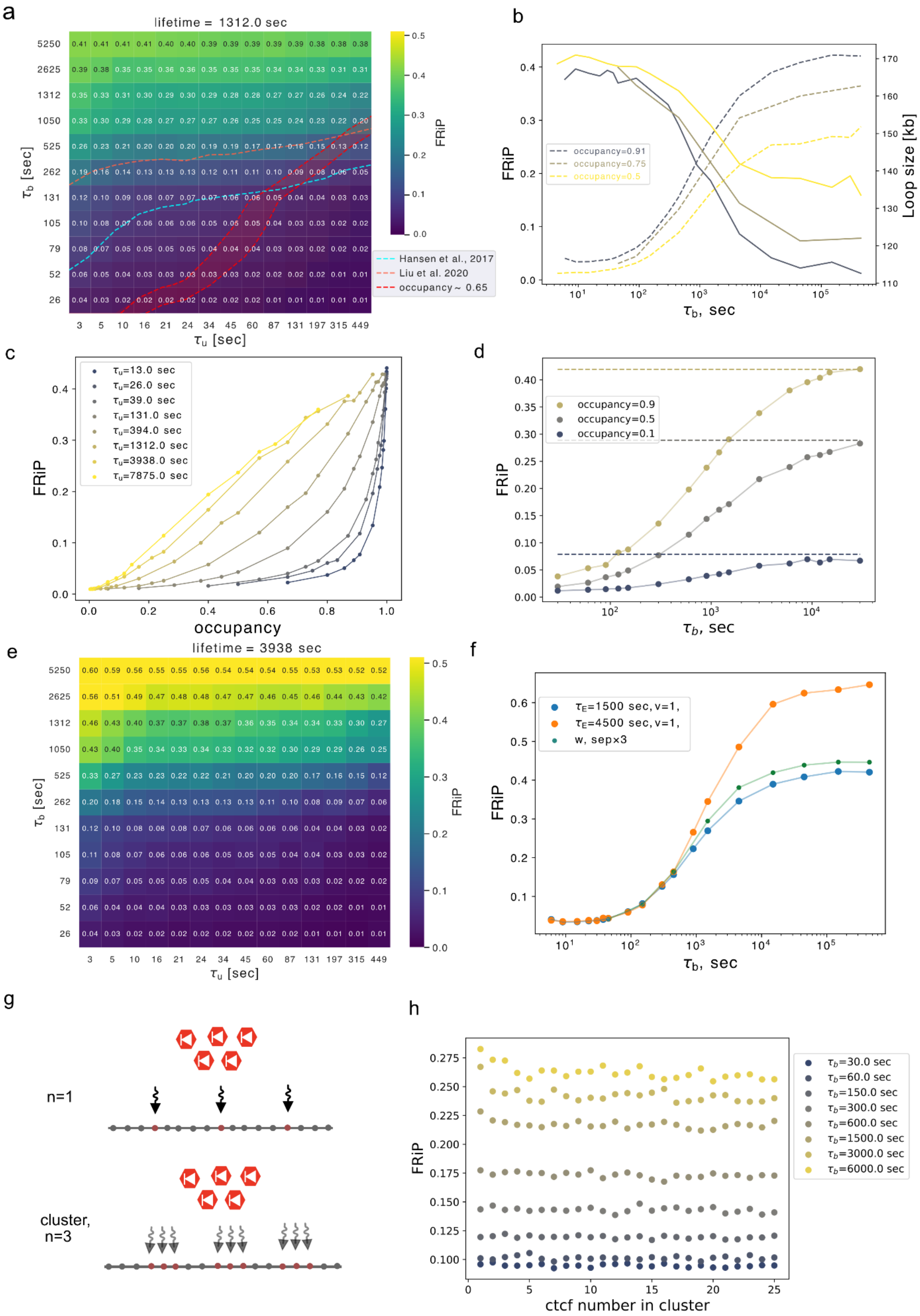
a
lifetime = 1312.0 sec
$\tau_b$ [sec]
$\tau_u$ [sec]
FRiP
Hansen et al., 2017
Liu et al. 2020
occupancy ~ 0.65
b
FRiP
Loop size [kb]
$\tau_b$, sec
occupancy=0.91
occupancy=0.75
occupancy=0.5
c
FRiP
occupancy
$\tau_u$=13.0 sec
$\tau_u$=26.0 sec
$\tau_u$=39.0 sec
$\tau_u$=131.0 sec
$\tau_u$=394.0 sec
$\tau_u$=1312.0 sec
$\tau_u$=3938.0 sec
$\tau_u$=7875.0 sec
d
FRiP
$\tau_b$, sec
occupancy=0.9
occupancy=0.5
occupancy=0.1
e
lifetime = 3938 sec
$\tau_b$ [sec]
$\tau_u$ [sec]
FRiP
f
FRiP
$\tau_b$, sec
$\tau_E$=1500 sec, v=1,
$\tau_E$=4500 sec, v=1,
w, sep×3
g
n=1
cluster, n=3
h
FRiP
ctcf number in cluster
$\tau_b$=30.0 sec
$\tau_b$=60.0 sec
$\tau_b$=150.0 sec
$\tau_b$=300.0 sec
$\tau_b$=600.0 sec
$\tau_b$=1500.0 sec
$\tau_b$=3000.0 sec
$\tau_b$=6000.0 sec

**Figure S3. FRiP dependency on extruder parameters and number of CTCF in cluster.**
Caption on next page.

**a.** Heatmaps of FRiP for $\boldsymbol{\tau}_b$ vs $\boldsymbol{\tau}_u$. Experimental values for FRiP are depicted with dashed contour lines, ranging from 0.07 (Hansen et al. 2017) to 0.16 (N. Q. Liu et al. 2021). Estimated experimental occupancy (0.7 (Sönmezer et al. 2021) ) is highlighted with red.
**b.** Inverse relationship between FRiP and loop size at multiple occupancies. Right y-axis indicates loop size (solid lines), the left y-axis indicates FRiP (dashed lines).
**c.** FRiP as a function of occupancy. FRiP increases linearly with occupancy at high $\boldsymbol{\tau}_u$, but increases sub-linearly when $\boldsymbol{\tau}_u \lesssim \boldsymbol{\tau}_E$ (=1312 sec)
**d.** FRiP approaches its value for static barriers (dashed lines) at large $\boldsymbol{\tau}_b$
**e.** heatmap of FRiP for increased extruder lifetime, as a function of bound and unbound time**.**
**f.** FRiP vs $\boldsymbol{\tau}_b$ for either increased lifetime or separation. While three-fold increased lifetime had strong effect at large $\boldsymbol{\tau}_b$ , three-fold increased separation had a weak effect.
**g.** Illustration of clusters of CTCF sites with a fixed average number of bound CTCFs at 50% occupancy. In this scenario each site has an increased unbound time (n*$\boldsymbol{\tau}_u$) and an occupancy 0.5/n.
**h.** FRiP as a function of CTCF number per cluster. To compute FRiP, we consider a fixed window (width=25) around the cluster of sites.

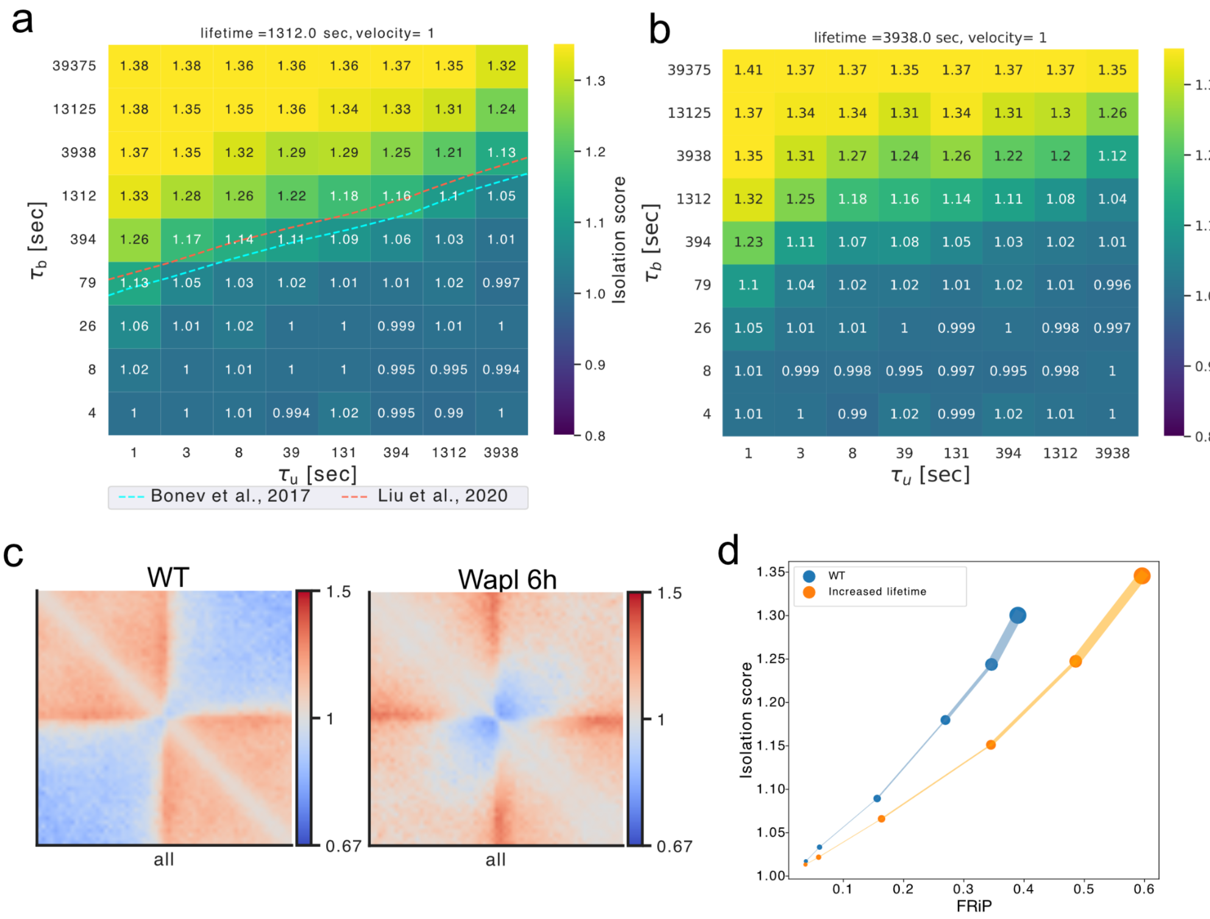


**Figure S4. Isolation strength of boundaries versus extruder lifetime.**

**a.** Heatmap of isolation score as a function of barrier lifetime and unbound time

**b.** Heatmap of isolation score for extruders with 3-times longer lifetimes than simulated WT

**c.** Average snippets around CTCF motifs overlap with CTCF chip-seq from Liu et al. 2020.

**d.** Isolation score versus FRiP. At a fixed extruder lifetime, isolation is correlated with FRiP. However, across extruder lifetimes, these scores are anti-correlated.

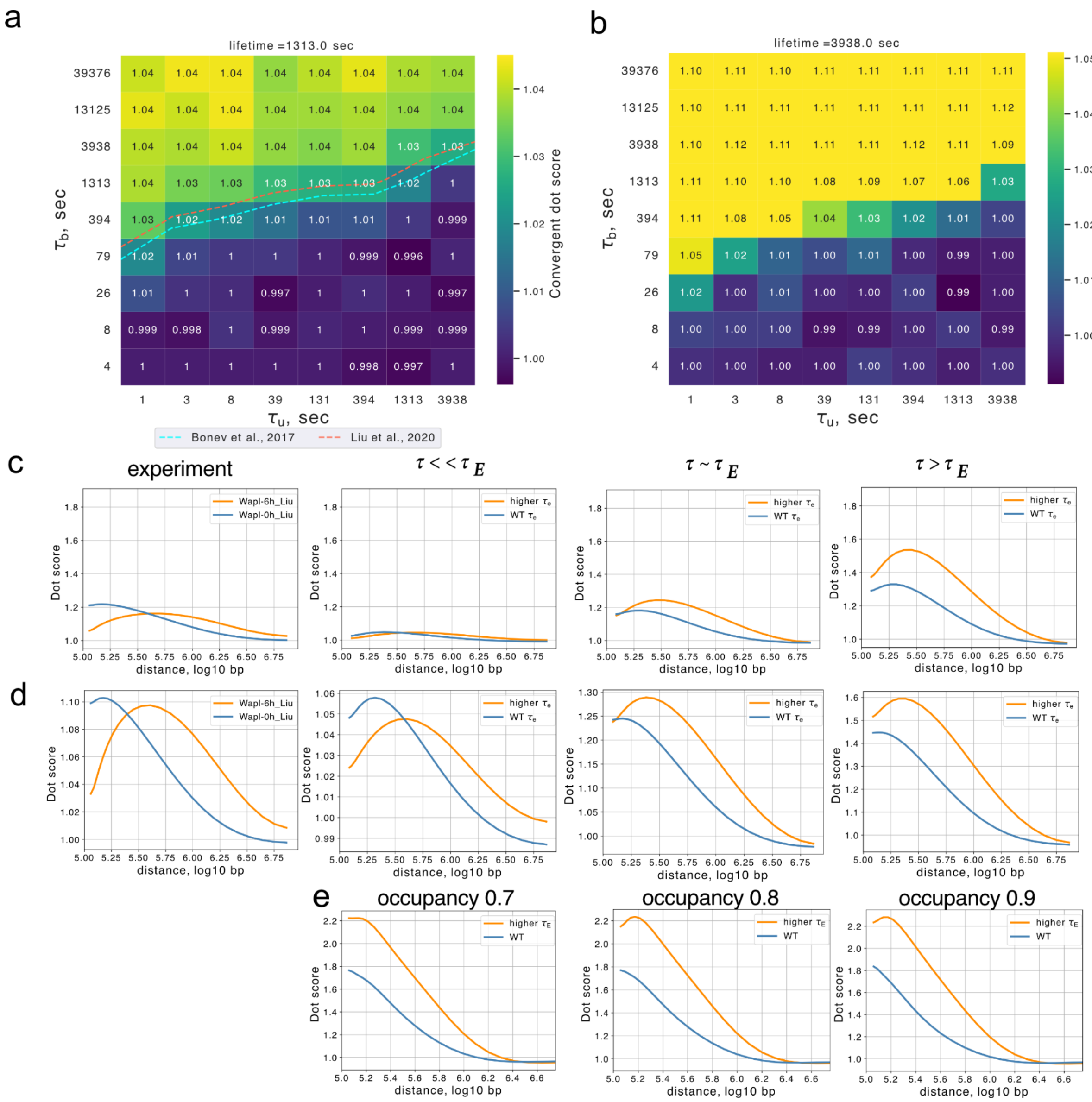


**Figure S5. Dot scores depend on extruder lifetime.**

**a.** Heatmap of dot score for barrier bound versus unbound times.

**b.** Heatmap at 3x higher lifetime.

**c.** A comparison of dot score magnitude between experimental data and simulation at various $\tau_b$ regimes, supporting the $\tau_b < \tau_E$ zone. Note the shared y-axis across all plots.

**d.** Overall dot score (i.e. averaged across all orientations) as a function of distance, for extruders with either WT (blue) and higher (orange) lifetimes at three CTCF bound times. Only simulations with $\tau_b<\tau_E$ displays similar behavior to experiments where the higher-lifetime curve starts lower and has a peak after the wildtype curve.

**e.** Dot score versus distance in simulations with static barriers is always higher for the higher lifetime extruders. This is similar to dynamic barriers with large $\tau_b$ ($\tau_b>>\tau_E$), yet differs from what is observed in experimental data.
